# Harnessing Ultrasound-Stimulated Phase Change Contrast Agents to Improve Antibiotic Efficacy Against Methicillin-Resistant *Staphylococcus aureus* Biofilms

**DOI:** 10.1101/2020.06.01.127340

**Authors:** Phillip G. Durham, Ashelyn E. Sidders, Paul A. Dayton, Brian P. Conlon, Virginie Papadopoulou, Sarah E. Rowe

**Affiliations:** Joint Department of Biomedical Engineering, The University of North Carolina and North Carolina State University, Chapel Hill, North Carolina 27599, USA; Eshelman School of Pharmacy, University of North Carolina, Chapel Hill, NC 27599, USA; Department of Microbiology and Immunology, University of North Carolina-Chapel Hill, Chapel Hill, North Carolina 27599, USA; Marsico Lung Institute, University of North Carolina at Chapel Hill, Chapel Hill, North Carolina 27599, USA

## Abstract

Bacterial biofilms, often associated with chronic infections, respond poorly to antibiotic therapy and frequently require surgical intervention. Biofilms harbor persister cells, metabolically indolent cells, which are tolerant to most conventional antibiotics. In addition, the biofilm matrix can act as a physical barrier, impeding diffusion of antibiotics. Novel therapeutic approaches frequently improve biofilm killing, but usually fail to achieve eradication. Failure to eradicate the biofilm leads to chronic and relapsing infection, associated with major financial healthcare costs and significant morbidity and mortality. We address this problem with a two-pronged strategy using 1) antibiotics that target persister cells and 2) ultrasound-stimulated phase-change contrast agents (US-PCCA), which improve antibiotic penetration.

We previously demonstrated that rhamnolipids, produced by *Pseudomonas aeruginosa*, could induce aminoglycoside uptake in gram-positive organisms, leading to persister cell death. We have also shown that US-PCCA can transiently disrupt biological barriers to improve penetration of therapeutic macromolecules. We hypothesized that combining antibiotics which target persister cells with US-PCCA to improve drug penetration could eradicate methicillin resistant *S. aureus* (MRSA) biofilms. Aminoglycosides alone or in combination with US-PCCA displayed limited efficacy against MRSA biofilms. In contrast, the anti-persister combination of rhamnolipids and aminoglycosides combined with US-PCCA dramatically reduced biofilm viability, frequently culminating in complete eradication of the biofilm. These data demonstrate that biofilm eradication can be achieved using a combined approach of improving drug penetration of therapeutics that target persister cells.

## Introduction

*S. aureus* is one of the most important human bacterial pathogens and in 2017 was the cause of 20,000 bacteremia deaths in the US alone^1^. Infections range from minor skin and soft tissue infections (SSTI), implanted device infections to more serious infections such as osteomyelitis, endocarditis and pneumonia ^2,3^. In addition to the high degree of mortality, chronic and relapsing *S. aureus* infections are common and associated with significant morbidity. This is due to frequent treatment failure of *S. aureus* infections. This is best illustrated by SSTIs, with some studies suggesting treatment failure rates as high as 45% and a recurrence rate of 70% ^4^. Importantly the failure of antibiotic therapy cannot be adequately explained by antibiotic resistance ^1^. Failure to clear the infection leads to a need for prolonged antibiotic therapies, increased morbidity and mortality, increased likelihood of antibiotic resistance development as well as an enormous financial healthcare burden.

*S. aureus* forms biofilms, bacterial cells embedded in a self-produced extracellular matrix, which act as a protective barrier from the host immune response and other environmental assaults. Biofilms expand up to 1200μm in thickness when attached to indwelling devices such as catheters ^5^. Non-surface attached biofilms, in chronic wounds and chronic lung infections, harbor smaller non-surface attached cell aggregates ranging from 2-200μm in diameter ^5,6^. These biofilm aggregates are often surrounded by inflammatory immune cells such as neutrophils and embedded in a secondary host produced matrix such as mucus, pus or wound slough ^7^. Consequently, biofilm-embedded cells have limited access to nutrients and oxygen and are coerced into a metabolically indolent state ^8^.

It has long been appreciated that biofilms respond poorly to antibiotics ^7, 9-12^. Most conventional bactericidal antibiotics kill by corrupting ATP-dependent cellular processes; aminoglycosides target translation, fluoroquinolones target DNA synthesis, rifampicin targets transcription and β-lactams and glycopeptides target cell wall synthesis ^13, 14^. Cells that survive lethal doses of antibiotics in the absence of a classical resistance mechanism are called antibiotic tolerant persister cells ^15^. Biofilms are made up of a high proportion of persister cells ^15- 18^. They are distinct from resistant cells as they cannot grow in the presence of the drug. However, once the drug is removed, persisters grow and repopulate a biofilm and cause a relapse in infection ^13^. Anti-persister antibiotics which kill independently of the metabolic state of the cell are more effective against biofilms than conventional antibiotics ^19-22^. Tobramycin, an aminoglycoside that requires active proton motive force (PMF) for uptake into the cell is inactive against non-respiring cells, anaerobically growing cells, small colony variants and metabolically inactive cells within a biofilm ^20^. We previously reported that rhamnolipids, biosurfactants produced by *P. aeruginosa*, permeabilize the *S. aureus* membrane to allow PMF-independent diffusion of tobramycin into the cell ^20,14, 22^. This combination of tobramycin and rhamnolipids (TOB/RL) rapidly sterilized in vitro planktonic cultures as well as non-respiring cells, anaerobically growing cells and small colony variants. However, despite this potent anti-persister activity, TOB/RL reduced biofilm viability by ∼3-logs but failed to achieve eradication ^20^. Notwithstanding the promise of this strategy, eradication of biofilms is arduous, even in vitro, indicating that factors other than the metabolic state of the biofilm-embedded cells are impeding therapy.

The biofilm matrix can act as a physical barrier to drug penetration. Penetration of oxacillin, vancomycin, cefotaxime, chloramphenicol and aminoglycoside antibiotics are impeded to some extent into *S. aureus* biofilms ^23-25^. Consequently, novel methods of drug delivery into biofilms is a growing area of interest. Ultrasound is a safe, commonplace, portable and relatively inexpensive modality typically used in medical imaging. This imaging capability has been expanded through the use of intravenously administered microbubbles as a contrast agent. These microbubbles are also used in a growing number of therapeutic applications to enhance biological effects, which include transdermal drug delivery ^26^ and transient permeabilization of the blood brain barrier ^27^.

When exposed to an ultrasound wave, gas-filled microbubbles in solution will oscillate, with the positive pressure cycle resulting in compression and the negative pressure cycle causing the bubble to expand. In an ultrasound field, microbubbles experience stable cavitation (continuous expansion and contraction) at lower pressures or inertial cavitation (violent collapse of the bubble) at higher pressures ^28^. Stable cavitation results in microstreaming; fluid movement around the bubble which induces shear stress to nearby structures (such as biofilms). At higher pressures, inertial cavitation can result in a shockwave, producing high temperatures at a small focus, and create microjets from the directional collapse of the bubble which can puncture host cells and disrupt physical barriers ^29^. Both of these pressure regimes have potential for therapeutic applications of ultrasound-mediated microbubble cavitation. Despite the potential of microbubbles to enhance drug delivery, their size (typically 1-4 micron in diameter) and short half-life once injected into solution may limit penetration and subsequent disruption of biofilms.

We hypothesized that phase change contrast agents (PCCA), submicron liquid particles (typically 100-400 nanometers in diameter) may be better equipped to penetrate a biofilm. PCCAs generally consist of a liquid perfluorocarbon droplet stabilized by a phospholipid shell. With appropriate ultrasound stimulation, PCCA can convert from the liquid phase to gas, generating a microbubble in their place (Fig. 1a). This process of “acoustic droplet vaporization” (ADV) may enhance drug penetration into biofilms as microbubbles over-expand before reaching their final diameter. Prior to activation, these particles are significantly more stable than microbubbles, with the potential to diffuse into biofilms due to their small size (Fig. 1b). Additionally, with continued ultrasound application, the resulting microbubbles can generate microstreaming, shear stress and microjets as they undergo cavitation (Fig. 1b). We hypothesized that PCCA, in combination with ultrasound (US-PCCA) and antibiotics that target persister cells is a novel biofilm eradication strategy.

**Fig 1.**
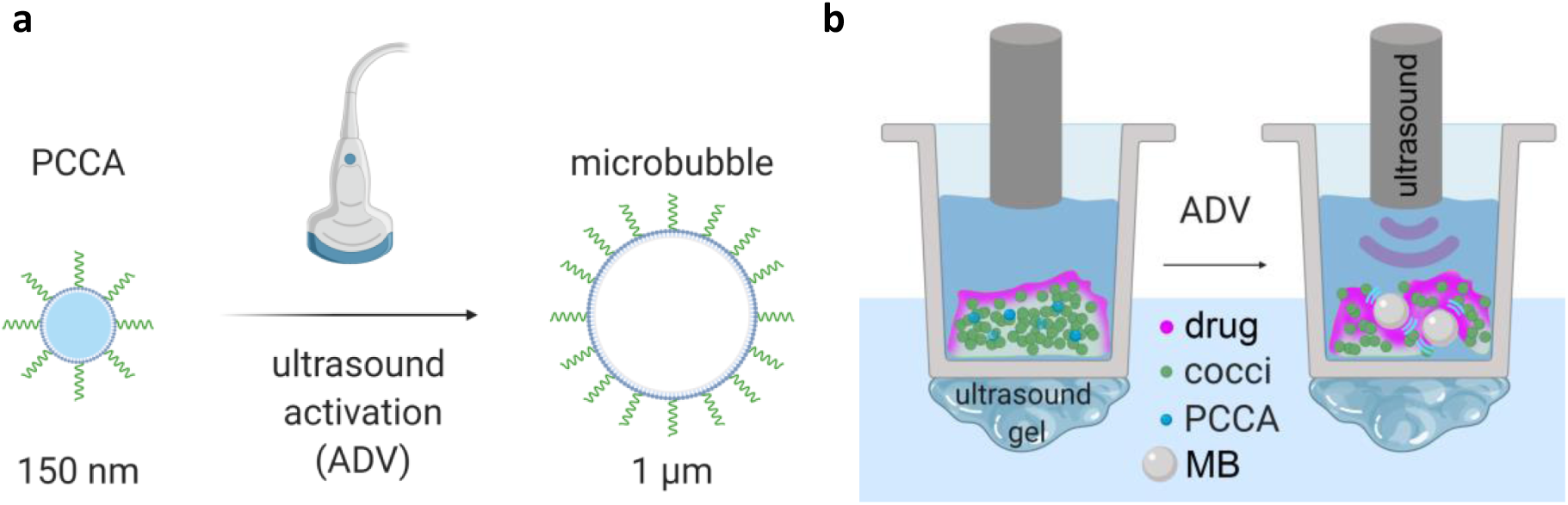
PCCA and ultrasound disrupts biofilm and increases drug penetration. Nanoscale PCCA are in a stable liquid phase. When exposed to ultrasound, the lipid shell containing superheated liquid perfluorocarbon is destabilized, causing the liquid to vaporize (acoustic droplet vaporization, ADV) to the gas phase and expand into a microbubble (a). The stability and small size of PCCA makes them ideal to diffuse into biofilms prior to ultrasound application. Ultrasound stimulation can vaporize PCCA to microbubbles that can physically disrupt biofilms and enhance drug penetration (b).

## Results

We first identified drugs with efficacy against biofilms. Antibiotics were chosen based on clinical relevance or previously reported anti-biofilm efficacy in vitro. Mature MRSA biofilms (USA300 LAC) were cultured for 24h in tissue culture treated plates before the addition of antibiotics. Following 24h of drug treatment, biofilms were washed and survivors were enumerated by plating. Tobramycin, mupirocin, vancomycin, and linezolid all caused a significant reduction in surviving biofilm cells (Fig. 2a). In contrast, levofloxacin and gentamicin showed no efficacy against biofilms at clinically achievable concentrations found in serum (C_max_) ^24, 25^ (Fig. 2a).

**Fig. 2:**
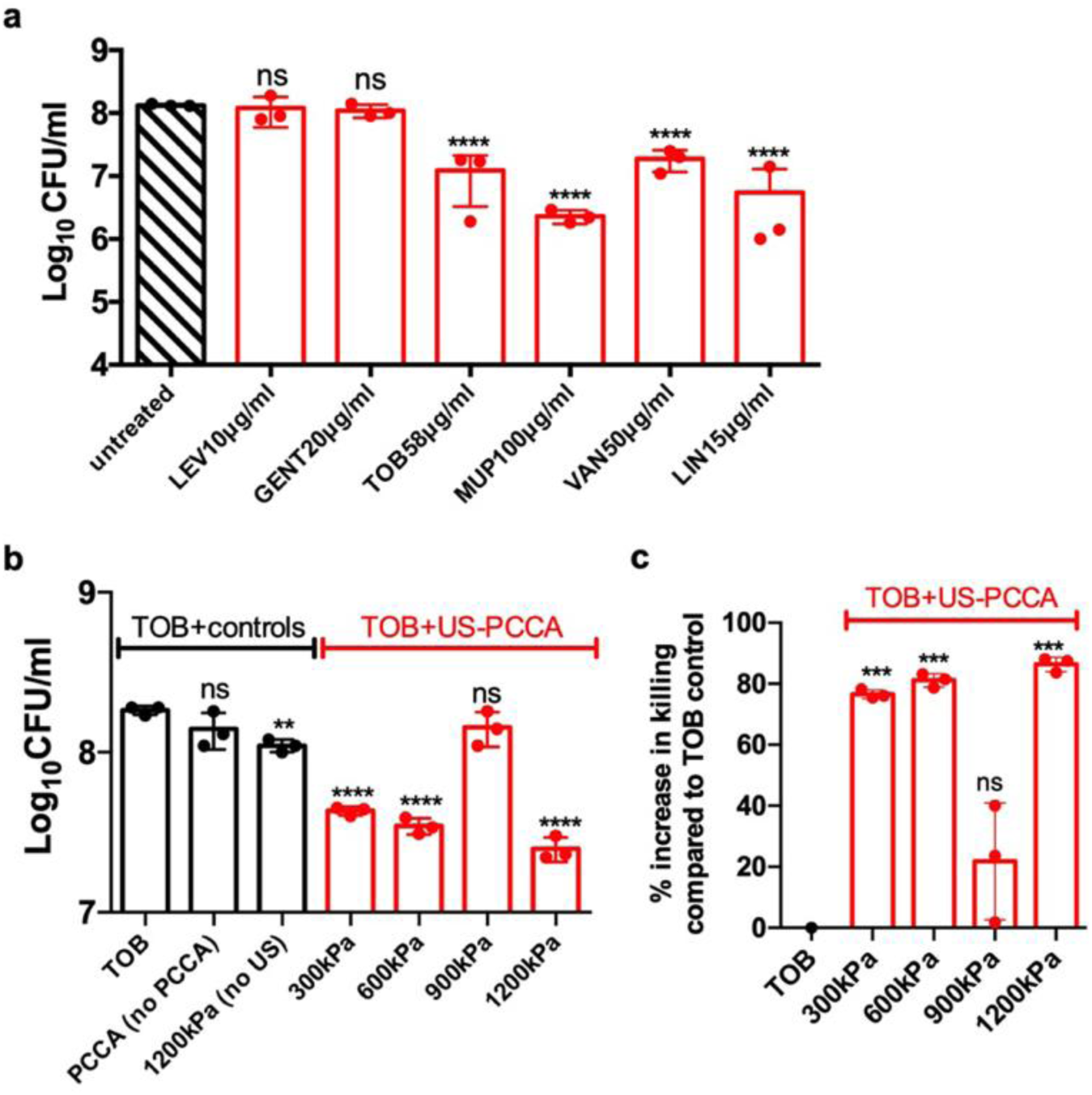
MRSA biofilms display high tolerance to clinically relevant antibiotics. MRSA strain LAC biofilms were cultured overnight in brain-heart infusion (BHI) media in 12-well tissue culture treated plates. Biofilms were washed and treated with antibiotics (a-c). Where indicated, 12-well plates were transferred to a custom-built temperature-controlled 37°C water bath alignment setup. PCCA were added and 30s ultrasound exposure was applied at indicated pressures and 20% duty cycle (b-c). After 24h, biofilms were washed, sonicated for disruption and surviving cells were enumerated by serial dilution plating. Survivors were presented as log_10_CFU/ml (a-b) or as % increase in killing compared to tobramycin control (c) (extrapolated from a). The averages of *n* = 3 biologically independent samples are shown. The error bars represent the standard deviation. Statistical significance was determined using a one-way analysis of variance (ANOVA) with Dunnett’s (a, c) or Sidak’s multiple comparison test (b). *, **, ** denotes P<0.005, P<0.0005, P<0.0001, respectively. LEV, levofloxacin; GENT, gentamicin; TOB, tobramycin; MUP, mupirocin; VAN, vancomycin; LIN, linezolid, ns, not significant; US-PCCA, ultrasound-stimulated phase change contrast agents.

Next, we tested the ability of 30 second (s) US-PCCA treatment to potentiate tobramycin efficacy. Previous studies have indicated that negatively charged components of the biofilm matrix such as extracellular DNA and certain components of polysaccharides impede penetration of positively charged aminoglycosides such as tobramycin ^25, 30, 31^. We hypothesized that US-PCCA might improve tobramycin penetration into biofilms and increase its efficacy. Mature biofilms were washed and transferred to a custom-built temperature-controlled 37°C water bath alignment setup. Tobramycin and PCCAs were added and ultrasound applied at a range of rarefactional pressures (300-1200kPa). We found that tobramycin efficacy was significantly enhanced at pressures of 300, 600 and 1200 but not 900KPa in the presence of PCCAs (Fig. 2b-c). We confirmed that the addition of PCCA in the absence of ultrasound had no impact on biofilm viability. Similarly, we anticipated that ultrasound alone, in the absence of PCCA would be ineffective, however 1200 kPa did cause a small but significant reduction in surviving cells in the absence of PCCA (Fig. 2b), indicating that potentiation seen at the highest pressure (1200kPa) may not be entirely attributable to PCCA activity. In order to investigate the potentiation effects of PCCA specifically in the regime below ultrasound-alone effects, the higher pressures (900 and 1200kPa) were not evaluated further and the duty cycle lowered to 10% for subsequent experiments. The lower pressures, 300 and 600kPa, in combination with PCCA were determined to be most effective at potentiating tobramycin efficacy. This is consistent with our previous findings where lower pressures (above the ADV threshold) resulted in more persistent cavitation activity during a 30s ultrasound exposure and was consistently greatest at macromolecule drug delivery across colorectal adenocarcinoma monolayers ^32^.

Next, we tested the ability of US-PCCA to potentiate mupirocin, vancomycin and linezolid/rifampicin. Mupirocin is a carboxylic acid topical antibiotic commonly used to treat *S. aureus* infections that binds to the isoleucyl-tRNA and prevents isoleucine incorporation into proteins ^33^. US-PCCA caused a very slight increase in mupirocin killing (41% increase in killing) that was statistically significant but of questionable biological significance (Fig. 2a-b).

Vancomycin is a glycopeptide that is the frontline antibiotic to treat MRSA infections. This antibiotic acts by binding to the D-Ala-D-ala residues of the membrane bound cell wall precursor, lipid II, preventing its incorporation and stalling active peptidoglycan synthesis ^34^. Importantly, some studies have indicated that vancomycin penetration is impeded into biofilms ^24^. US-PCCA potentiated vancomycin killing of biofilm-associated cells by 93% (Fig. 3a-b), likely by improving penetration. Notably, potentiation of vancomycin was seen with the C_max_ ^35^ indicating that at a clinically relevant concentration, US-PCCA has the capacity to improve biofilm killing of the front-line antibiotic used to treat MRSA infections.

**Fig. 3:**
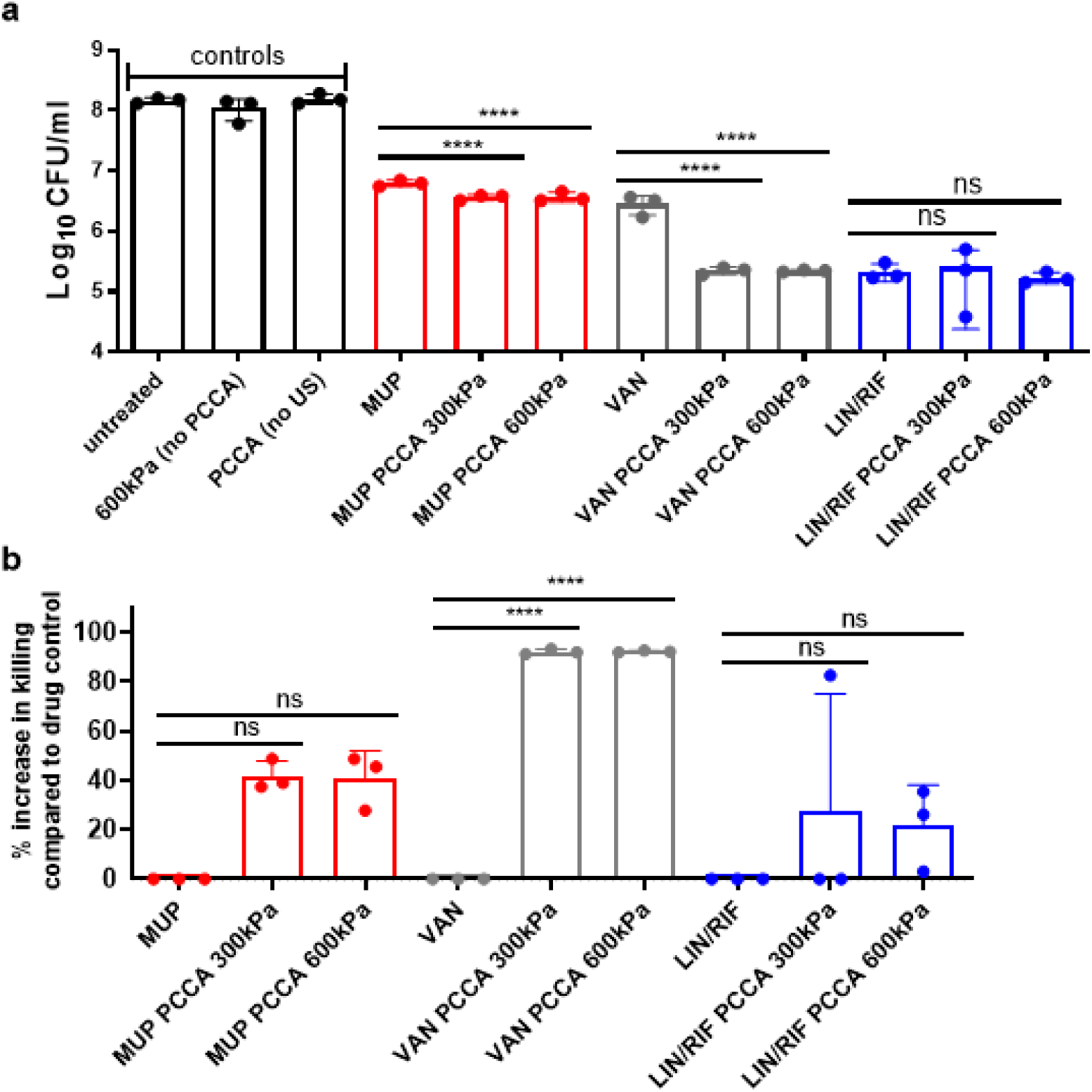
US-PCCA improve vancomycin killing of MRSA biofilms. MRSA strain LAC biofilms were cultured overnight in brain-heart infusion (BHI) media in 24-well tissue culture treated plates. Biofilms were washed and treated with antibiotics and transferred to a custom-built temperature-controlled 37°C water bath alignment setup. PCCA were added and 30s ultrasound exposure was applied at indicated pressures and 10% duty cycle. After 24h, biofilms were washed, sonicated for disruption and surviving cells were enumerated by serial dilution plating. Survivors were presented as log_10_CFU/ml (a) or as % increase in killing compared to antibiotic control (b) (extrapolated from a). The averages of *n* = 3 biologically independent samples are shown. The error bars represent the standard deviation. Statistical significance was determined using a one-way analysis of variance (ANOVA) with Sidak’s multiple comparison test. ** denotes P<0.0001, respectively. MUP, 100μg/ml mupirocin; VAN, 50μg/ml vancomycin; LIN, 15μg/ml linezolid; RIF, 10μg/ml rifampicin; ns, not significant; US-PCCA, ultrasound-stimulated phase change contrast agents.

Linezolid is an oxazolidinone protein synthesis inhibitor that is sometimes combined with the transcriptional inhibitor, rifampicin, for the treatment of *S. aureus* infections ^36, 37^. Linezolid/rifampicin reduced viable cells within the biofilm by almost 3-logs but was not significantly potentiated by US-PCCA. This suggests that US-PCCA has the ability to potentiate some conventional antibiotics but not others. It is possible that US-PCCA does not potentiate the killing of mupirocin and linezolid/rifampicin because the penetration of these drugs is not impeded into biofilms.

Although the increased killing of biofilm-associated cells with conventional antibiotics shows promise, we hypothesized that regardless of penetration, antibiotic tolerant persister cells in the biofilm are surviving and thus impeding biofilm eradication. We predicted that utilizing US-PCCA to increase penetration of drugs active against antibiotic tolerant persister cells could achieve eradication of a biofilm.

Daptomycin is a lipopeptide antibiotic which inserts into the cell membrane and disrupts fluid membrane microdomains ^38^. Daptomycin has potent activity against recalcitrant populations of *S. aureus*, including biofilms ^39, 40^. Daptomycin in combination with linezolid (DAP/LIN) is the treatment recommended for persistent MRSA bacteremia or vancomycin failure in the Infectious Diseases Society of America 2011 MRSA treatment guidelines ^41^. We found that US-PCCA increased DAP/LIN killing of MRSA biofilms by 83% and 90% at 300kPa and 600kPa, respectively (Fig. 4a-b).

**Fig. 4:**
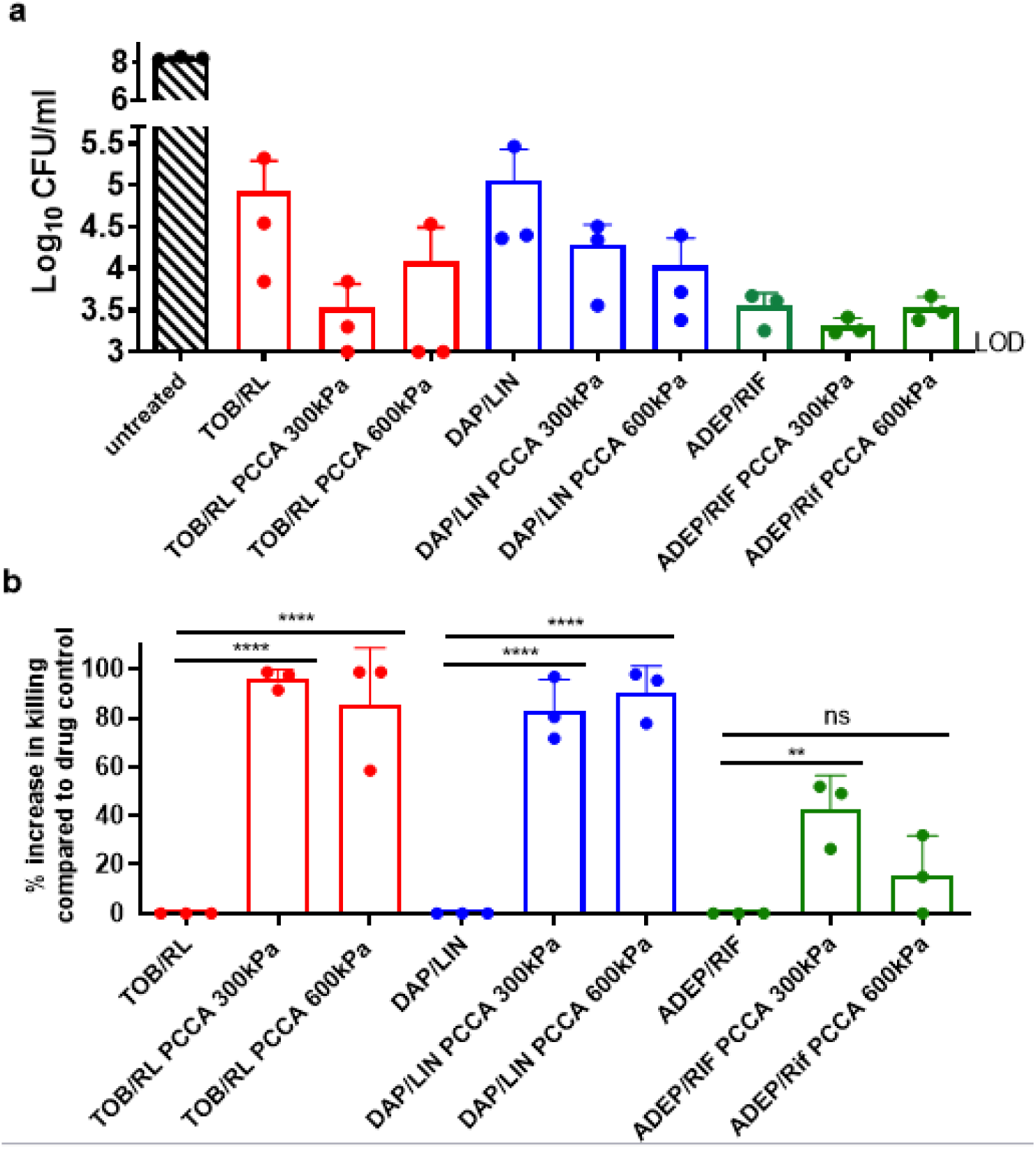
US-PCCA improves anti-persister antibiotic therapy against MRSA biofilms. MRSA strain LAC biofilms were cultured overnight in brain-heart infusion (BHI) media in 12-well (TOB/RL) or 24-well (all other drugs) tissue culture treated plates. Biofilms were washed and treated with antibiotics and transferred to a custom-built temperature-controlled 37°C water bath alignment setup. PCCAs were added and 30s ultrasound exposure was applied at indicated pressures and 10% duty cycle. After 24h, biofilms were washed, sonicated for disruption and surviving cells were enumerated by serial dilution plating. Survivors were presented as log_10_CFU/ml (a) or as % increase in killing compared to antibiotic control (b) (extrapolated from a). The averages of *n* = 3 biologically independent samples are shown. The error bars represent the standard deviation. Statistical significance was determined using a one-way analysis of variance (ANOVA) with Sidak’s multiple comparison test (b). *, ** denotes P<0.005, P<0.0001, respectively. TOB, 58μg/ml tobramycin; RL, 30μg/ml rhamnolipids; DAP, 100μg/ml daptomycin; LIN, 15μg/ml linezolid; RIF, 10μg/ml rifampicin; ADEP, 5μg/ml acyldepsipeptide; ns, not significant; US-PCCA, ultrasound-stimulated phase change contrast agents, LOD; limit of detection.

Next, we wanted to investigate if US-PCCA could improve efficacy of other drugs with anti-persister activity. Acyldepsipeptides (ADEPs) are activators of the ClpP protease. We previously reported that ADEPs sterilize persisters by activating the ClpP protease and causing the cell to self-digest in an ATP-independent manner ^19^. ADEP in combination with rifampicin reduced biofilm cells by >4-logs in 24h. US-PCCA significantly potentiated efficacy of ADEP/RIF at 300kPa but not 600kPa (Fig. 4a-b).

Tobramycin combined with rhamnolipids (TOB/RL), has potent anti-persister activity and has eradiated several recalcitrant populations including non-respiring cells, anaerobically growing cells and small colony variants ^20^. Despite this potent anti-persister activity, TOB/RL only reduced biofilm viability by ∼3-logs and failed to achieve biofilm eradication ^20^. We reasoned that drug penetration might be inhibited into the biofilms and hypothesized that improving penetration could lead to biofilm eradication. Applying US-PCCA in combination with TOB/RL increased killing of biofilm cells by 96% and 85% at 300kPa and 600kPa, respectively (Fig. 4a-b). Importantly, half of the biological replicates were eradicated to the limit of detection. Together this data indicates that anti-persister drugs have potent anti-biofilm activity and this can be potentiated further by improving penetration using US-PCCA.

## Discussion

*S. aureus* biofilms rarely resolve with antibiotic treatment alone and usually require surgical intervention (debridement, drainage, incision) ^42^. Many antibiotics reduce bacterial burdens within biofilms but eradication represents an arduous challenge even in vitro ^5, 15^. In this study, we combine two anti-biofilm strategies to achieve eradication (Fig. 5). Biofilm killing by conventional antibiotics with impeded penetration is improved by US-PCCA (Fig.2b-c, Fig.3a-b, Fig. 5), highlighting the therapeutic potential despite falling short of eradication. Targeting biofilms with anti-persister drugs increases efficacy compared to conventional antibiotics but fails to eradicate (Fig. 4a). Combining these approaches greatly improves biofilm killing, and in some cases resulted in eradication of the biofilm (Fig. 4a, Fig. 5).

**Fig. 5:**
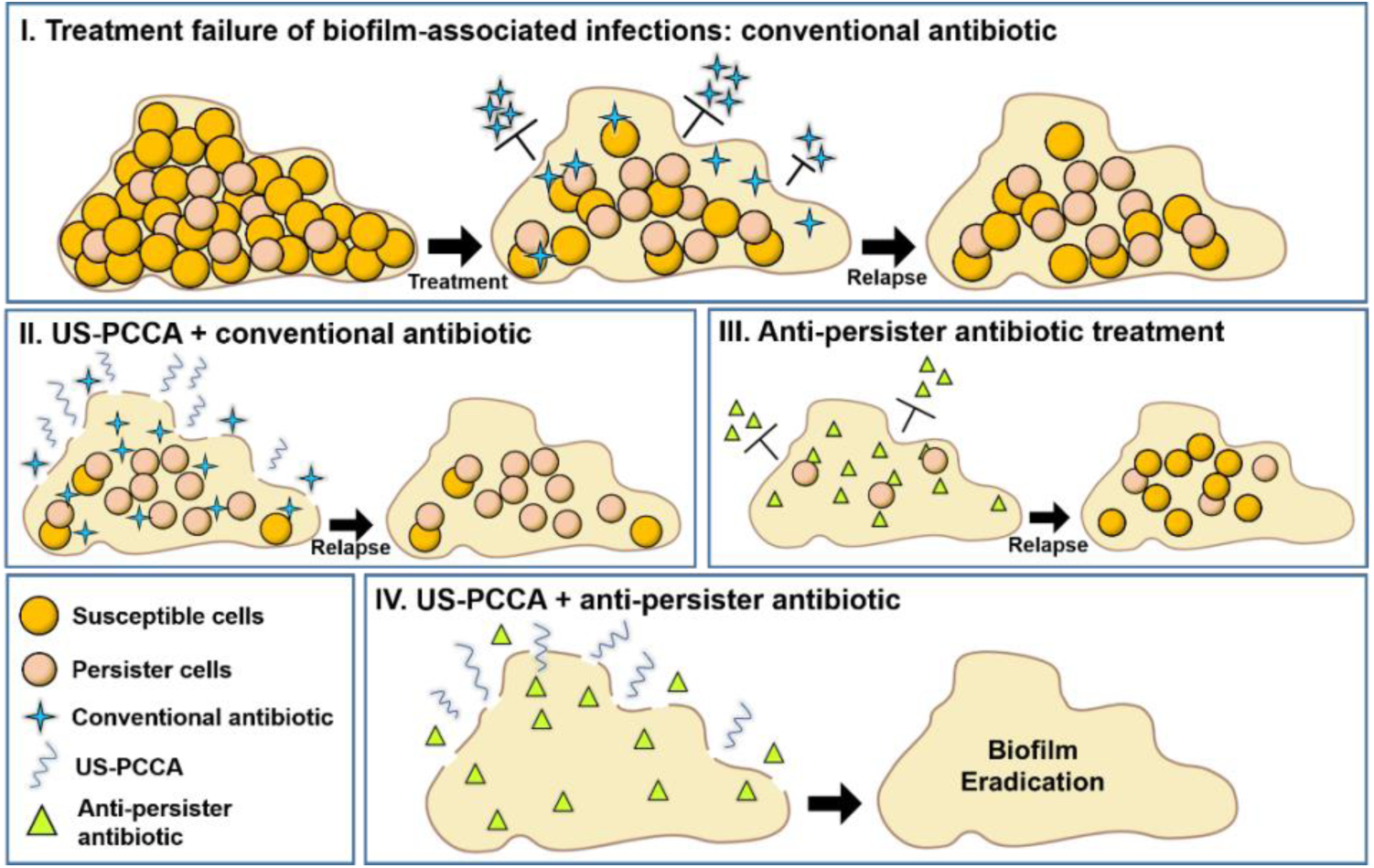
Schematic representing a dual approach to improving antibiotic therapy against *S. aureus* biofilms. (I) Biofilms display remarkable tolerance to antibiotics. Susceptible cells at the biofilm periphery die (susceptible cells) while less metabolically active cells within the biofilm are tolerant to conventional antibiotics (persister cells). Failure to eradicate the biofilm leads to relapse in infection following removal of the antibiotic. (II) Improving penetration of conventional antibiotics using US-PCCA will improve efficacy of some conventional antibiotics that do not penetrate well through the biofilm matrix. This strategy is futile as it does not improve killing of persister cells and leads to relapse. (III) Targeting biofilms with antibiotics which kill persister cells (anti-persister antibiotics) improves efficacy but if drug penetration is impeded into the biofilm, some persister cells will remain following drug treatment and lead to relapse. (IV) Improving penetration of anti-persister antibiotics into the biofilm can lead to the eradication of all cells within a biofilm and prevent relapse of infection following removal of the antibiotic.

Antibiotic treatment failure is a complex issue that imposes a heavy burden on global public health. The last new class of antibiotics to be approved by the FDA was in 2003 ^43^. Unlike drugs for chronic illnesses that are administered for life (e.g. heart disease, diabetes), antibiotic regimens are comparatively short, rendering the profitability of antibiotic development low ^44^. The void in the drug discovery pipeline makes sensitizing recalcitrant bacterial populations to already approved therapeutics a promising approach. The use of ultrasound and cavitation-enhancing agents for antibacterial applications, recently termed “sonobactericide”, was first published in 2011 ^45^. While the field is still developing, a significant prospect of therapeutic ultrasound as a mechanical approach to enhance drug efficacy is its compatibility with any molecular therapeutic.

Microbubble oscillation has been shown to cause discrete morphologic changes in a *P. aeruginosa* biofilm ^46^. Disruption of the physical structure of the biofilm may increase penetration depth of molecules which would otherwise be impeded. Disruption of the biofilm may have other indirect effects on drug efficacy. For example, bacterial biofilms are often hypoxic due to the diffusional distance limit of oxygen. Creating holes in the biofilm may allow oxygen penetration and stimulate the metabolic state of the residing persister cells, rendering them sensitive to antibiotics. In support of this, ultrasound in combination with microbubbles has previously been reported to alter the metabolic state of bacterial biofilms ^46, 47^.

We hypothesized that PCCAs may be more efficient than microbubbles at penetrating biofilms due to their relatively small size and increased stability. The use of US-PCCAs has previously shown to increase vancomycin killing of MRSA biofilms ^48^. In contrast to the current study, Hu et al. used perfluoropentane as the perfluorocarbon core, which requires higher pressures than octofluoropropane to vaporize. Even in the absence of an antibiotic, US-PCCA caused a significant reduction in biofilm matrix and metabolic activity measured by three-dimensional fluorescence imaging and resazurin ^47^. The difference in quantification method makes comparison with the previous study difficult (we enumerated bacterial survivors), however our results demonstrate a significant improvement in efficacy using shorter treatment times (30 s vs 5 minutes)^49^. In addition, the low boiling point PCCAs used in the current study present the advantage that the same low-pressure ultrasound settings can be used for both ADV and subsequent microbubble cavitation. Indeed, this can be achieved with clinically available ultrasound hardware at pressures below the FDA set limits for diagnostic imaging. Additionally, PCCA formulation is a variant of FDA-approved ultrasound contrast microbubbles that have been clinically used for over 25 years in Europe, Asia and USA. This approach may improve the efficacy of existing approved drugs without the additional need for the extensive regulatory approval which accompanies a new molecule. Likewise, as it uses ultrasound parameters that are achievable with clinically available equipment, this has the potential for rapid translation to clinical practice without the need for further technological development.

The ultrasound parameters used in our study mostly varied acoustic pressure and have not yet been optimized for in vivo application. While acoustic pressure is a large contributor to PCCA activation and stimulation, other parameters of frequency, duty cycle, treatment time and PCCA concentration could be further evaluated. Ultrasound is also used clinically for debridement of wounds to disperse biofilms ^50^. Evaluation of PCCA drug potentiation using lower frequencies and higher intensities typical for this application could give further insight into clinical integration strategies. Future experiments will evaluate the potentiation of antibiotics in a *S. aureus* mouse skin and soft tissue infection (SSTI). For topical applications such as soft tissue infections, we believe maintaining cavitation activity for the duration of the treatment will be crucial for efficacy, as no new cavitation nuclei will be introduced as would be the case in intravenously administered PCCA (replenished by blood flow).

## Methods

### Biofilm assays

Biofilm assays were performed using the USA300 MRSA strain LAC. It is a highly characterized community-acquired MRSA (CA-MRSA) strain isolated in 2002 from an abscess of an inmate in Los Angeles County jail in California ^51^. LAC was cultured overnight (18h) in brain heart infusion (BHI) media (Oxoid) in biological triplicates. Each culture was diluted 1:150 in fresh media and 2-3ml was added to the wells of 24-well or 12-well tissue culture treated plates (Costar), respectively. Biofilms were covered with Breathe-Easier sealing strips (Sigma) and incubated at 37°C for 24h. Biofilms were carefully washed twice with PBS and fresh BHI media containing antibiotics was added. Biofilms were covered and incubated at 37°C for 24h. Biofilms were carefully washed twice with PBS before dispersal in a sonicating water bath (5min) and vigorous pipetting. Surviving cells were enumerated by serial dilution and plating. Antibiotics were added at concentrations similar to the C_max_ in humans; 10μg/ml levofloxacin^52^ (Alfa Aesar), 20μg/ml gentamicin^53^ (Fisher BioReagents), 58μg/ml tobramycin^54^ (Sigma), 50μg/ml vancomycin hydrochloride^35^ (MP Biomedicals), 15μg/ml linezolid^55^ (Cayman Chemical), 10μg/ml rifampicin^56^ (Fisher BioReagents), 100μg/ml daptomycin^57^ (Arcos Organics), with the exception of the topical antibiotic mupirocin (Sigma) (administered at 100μg/ml) and acyldepsipeptide antibiotic (ADEP4) which was added at 10x MIC (10μg/ml) which previously showed efficacy against *S. aureus* biofilms^19^. For daptomycin activity, the media was supplemented with 50mg/L of Ca^2+^ ions. Where indicated tobramycin was supplemented with 30μg/ml rhamnolipids^22^ (50/50 mix of mono- and di-rhamnolipids, Sigma). Where indicated biofilms were treated with PCCA and ultrasound.

### PCCA Generation

Phase change contrast agents were generated as previously reported ^58^ [Sheeran et al. 2012]. Briefly, 1,2-distearoyl-sn-glycero-3-phosphocholine (DSPC) and 1,2-distearoyl-sn-glycero-3-phosphoethanolamine-N-methoxy(polyethylene-glycol)-2000 (DSPE-PEG2000) (Avanti Polar Lipids, Alabaster, AL, USA) were dissolved in 5% glycerol, 15% propylene glycol (both from Fisher Chemical, Waltham, MA, USA) in PBS (v/v) at a 1:9 ratio, to a total lipid concentration of 1 mg/ml. Lipid solution (1.5ml) was dispensed into 3ml crimp-top vials and degassed under vacuum for 30 minutes and then backfilled with octofluoropropane (OFP) gas (Fluoro Med, Round Rock, TX, USA). The vials were activated by mechanical agitation (VialMix, Bristol-Myers-Squibb, New York, NY, USA) to generate micron scale OFP bubbles with a lipid coat. The vials containing bubbles were cooled in an ethanol bath to -11C. Pressurized nitrogen (45 PSI) was introduced by piercing the septa with a needle and used to condense the gaseous octofluoropropane into a liquid, creating lipid-shelled perfluorocarbon submicron droplets (PCCA). Particle size and concentration was characterized the Accusizer Nano FX (Entegris, Billerica, MA, USA).

### Ultrasound Experiments

Ultrasound experiments were conducted in 12 or 24 well tissue culture plates using a custom fabricated water bath ultrasound alignment setup to maintain 37C during the experiment, similar to a design used previously with cell monolayers ^32^. Briefly, alignment guides were positioned above the wells to ensure reproducible transducer placement to the center of each well on top of the biofilm and 10mm from their bottom. To limit acoustic reflections and standing waves from the bottom of the well plate, ultrasound gel was applied on the outside bottom of each well before the plate was positioned in the water bath, the bottom of which was lined with acoustic absorber material. The water temperature was maintained at 37C throughout the experiment by placing the water bath setup on a heated plate and monitored by thermocouple. A 1.0MHz unfocused transducer (IP0102HP, Valpey Fisher Corp) was characterized via needle hydrophone and driven with an amplified 20- or 40-cycle sinusoidal signal defined on an arbitrary function generator (AFG3021C, Tektronix, Inc.; 3100LA Power Amplifier, ENI) at a pulse-repetition frequency of 5000 Hz (10% or 20% duty cycle). Peak negative pressures of 300, 600, 900 and 1200kPa were used in the experiments. To avoid ultrasound-alone effects on the biofilm, we focused on the lower pressures, 300 and 600kPa, determined most effective at potentiating tobramycin efficacy with PCCA and lowered the duty cycle from 20% in Fig. 1 to 10% in Fig. 2-3 as this was shown to have a more modest effect in our prior work and resulted in significant drug delivery ^32^. Where indicated, 10µl of PCCA was added to each well ((1.17 ± 0.4) × 10^11^ particles / mL, 0.18 μm diameter) and mixed gently by pipetting. The transducer was positioned in the well in the media above the biofilm and ultrasound treatment was applied for 30 seconds. Following treatment, each plate was incubated at 37C for 24h before enumerating survivors (described in detail above).

### Statistical information

The averages of *n* = 3 biologically independent samples are shown. The error bars represent the standard deviation of the mean. Statistical analysis was performed using Prism 8 (GraphPad) software. One-way ANOVA with Sidak’s or Dunnett’s multiple comparison test (as indicated in the figure legends). Statistical significance was defined as P < 0.05.

## Acknowledgements

This work was supported in part by NIH grants R01AI137273 and Cystic Fibrosis Research Grant to B.P.C and R33CA206939 to P.A.D. We are grateful to Nikki J. Wagner, Lauren C. Radlinski and Matthew Wolfgang for technical advice, as well as Brian Velasco for assistance with PCCA preparation.

